# Robust de novo design of protein binding proteins from target structural information alone

**DOI:** 10.1101/2021.09.04.459002

**Authors:** Longxing Cao, Brian Coventry, Inna Goreshnik, Buwei Huang, Joon Sung Park, Kevin M. Jude, Iva Marković, Rameshwar U. Kadam, Koen H.G. Verschueren, Kenneth Verstraete, Scott Thomas Russell Walsh, Nathaniel Bennett, Ashish Phal, Aerin Yang, Lisa Kozodoy, Michelle DeWitt, Lora Picton, Lauren Miller, Eva-Maria Strauch, Samer Halabiya, Bradley Hammerson, Wei Yang, Steffen Benard, Lance Stewart, Ian A. Wilson, Hannele Ruohola-Baker, Joseph Schlessinger, Sangwon Lee, Savvas N. Savvides, K. Christopher Garcia, David Baker

## Abstract

The design of proteins that bind to a specific site on the surface of a target protein using no information other than the three-dimensional structure of the target remains an outstanding challenge. We describe a general solution to this problem which starts with a broad exploration of the very large space of possible binding modes and interactions, and then intensifies the search in the most promising regions. We demonstrate its very broad applicability by de novo design of binding proteins to 12 diverse protein targets with very different shapes and surface properties. Biophysical characterization shows that the binders, which are all smaller than 65 amino acids, are hyperstable and bind their targets with nanomolar to picomolar affinities. We succeeded in solving crystal structures of four of the binder-target complexes, and all four are very close to the corresponding computational design models. Experimental data on nearly half a million computational designs and hundreds of thousands of point mutants provide detailed feedback on the strengths and limitations of the method and of our current understanding of protein-protein interactions, and should guide improvement of both. Our approach now enables targeted design of binders to sites of interest on a wide variety of proteins for therapeutic and diagnostic applications.

## Introduction

Protein interactions play critical roles in biology, and general approaches to disrupt or modulate these with designed proteins would have huge impact. While empirical laboratory selection approaches starting from very large antibody, DARPIN or other protein scaffold libraries can generate binders to protein targets, it is difficult at the outset to target a specific region on a target protein surface, and to sample the full space of possible binding modes. Computational methods can target specific target surface locations and provide a more principled and potentially much faster approach to binder generation than random library selection methods, as well as insight into the fundamental properties of protein interfaces (which must be understood for design to be successful). Most current methods for computationally designing proteins to bind to a target surface utilize information derived from native complex structures on specific sidechain interactions or protein backbone placements optimal for binding^1–3^. Computational docking of antibody scaffolds with varied loop geometries has yielded binders, but the designed binding modes have rarely been validated with high-resolution structures^4^. Binders have been generated starting from several computationally identified hot-spot residues, which were then used to guide the positioning of naturally occurring protein scaffolds^5^. However, for many target proteins, there are no obvious pockets or clefts on the protein surface into which a small number of privileged sidechains can be placed, and guidance by only a small number of hotspot residues limits the approach to a small fraction of possible interaction modes.

## Design Method

We sought to develop a general approach to design of high affinity binders to arbitrary protein targets that addresses two major challenges. First, in the general case, there are no clear sidechain interactions or secondary structure packing arrangements that can mediate strong interactions with the target; instead there are a very large number of individually very weak possible interactions. Second, the number of ways of choosing from these numerous weak interactions to incorporate into a single binding protein is combinatorially large, and any given protein backbone is unlikely to be able to simultaneously present sidechains that can encompass any preselected subset of these interactions. To motivate our approach, consider the simple analogy of a very difficult climbing wall with only a few good footholds or handholds distant from each other. Previous “hotspot” based approaches correspond to focusing on routes involving these footholds/handholds, but this greatly limits the possibilities and there may be no way to connect them into a successful route. An alternative is to first, identify all possible handholds and footholds, no matter how poor, second, have thousands of climbers select subsets of these, and try to climb the wall, third, identify those routes that were most promising, and fourth, have a second group of climbers explore them in detail. Following this analogy, we devised a multi-step approach to overcome the above two challenges by 1) enumerating a large and comprehensive set of disembodied sidechain interactions with the target surface, 2) identifying from large in silico libraries of protein backbones those that can host many of these sidechains without clashing with the target, 3) identifying recurrent backbone motifs in these structures, and 4) generating and placing against the target a second round of scaffolds containing these interacting motifs (**Fig. 1a**). Steps 1 and 2 search the space very widely, while steps 3 and 4 intensify search in the most promising regions. We describe and motivate each step in the following paragraphs.

**Figure 1:**
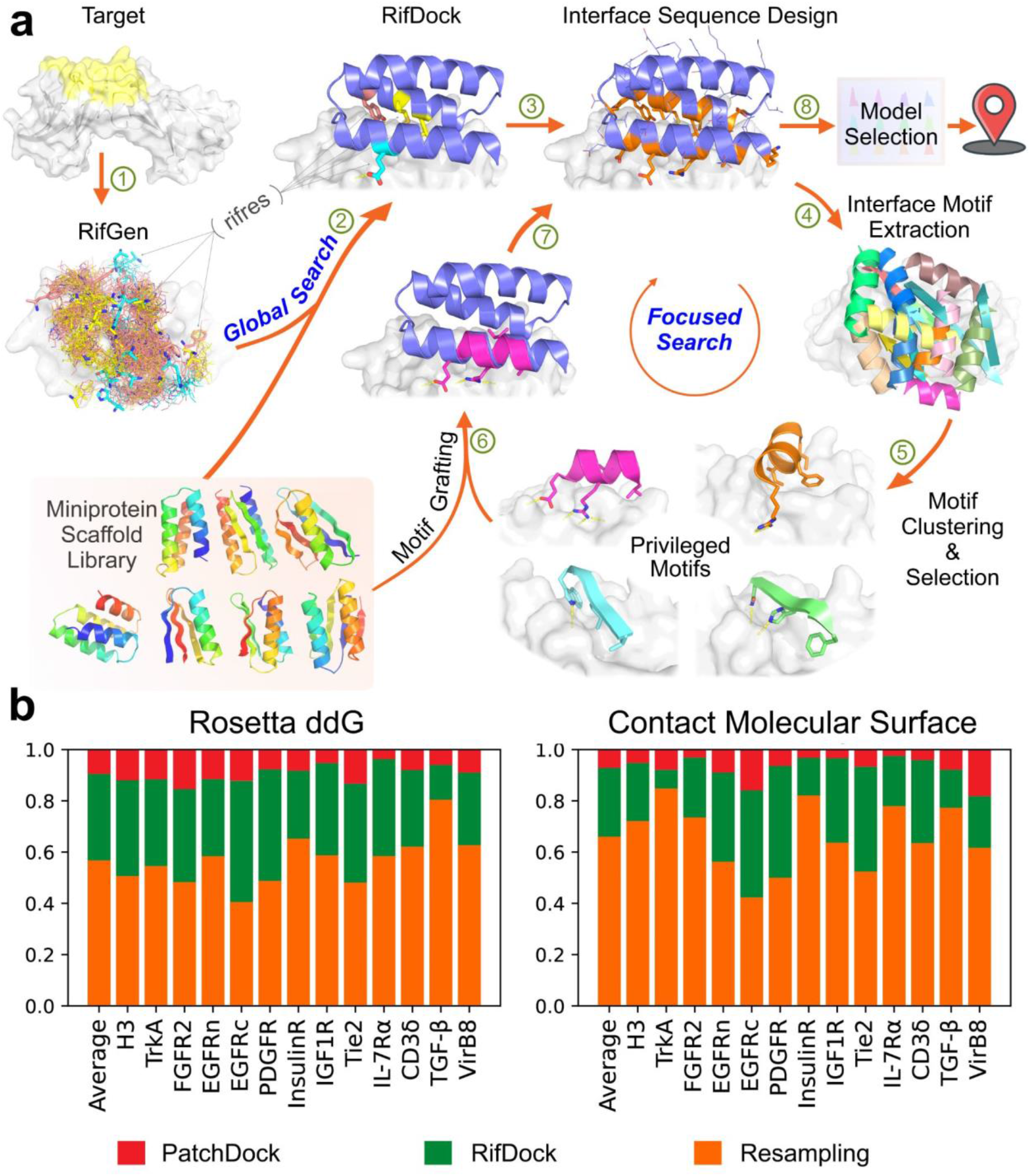
Overview of the de novo protein binder design pipeline. **a**, Schematic of our two stage binder design approach. In the global search stage, billions of disembodied amino acids are docked onto the selected targeting region and the positioning of the scaffolds is guided by the favorable sidechain interactions. The interface sequences are then designed to maximize interaction with the target. In the focused search stage, the interface motifs are extracted, clustered. The privileged motifs are then selected to guide another round of docking and design. Designs are then selected for experimental characterization based on computational metrics. **b**, Comparison of sampling efficiency of PatchDock, RifDock, and resampling protocols. Bar graph shows the distribution over the three approaches of the top 1% of binders based on Rosetta ddg and contact molecular surface after pooling equal-CPU-time dock-and-design trajectories for each of the 13 target sites and averaging per-target distributions (see Methods).

We began by docking disembodied amino acids against the target protein, and storing the backbone coordinates and target binding energies of the typically billions of amino acids making favorable hydrogen bonding or non-polar interactions in a 6-dimensional spatial hash table for rapid lookup (**Fig. 1a;** see methods). This “rotamer interaction field” (RIF) enables rapid approximation of the target interaction energy achievable by a protein scaffold docked against a target based on its backbone coordinates alone (with no need for time consuming sidechain sampling)--for each dock, the target interaction energies of each of the matching amino acids in the hash table are summed. A related approach was used for small molecule binder design^6^; since protein targets are so much bigger, and non-polar interactions are the primary driving force for protein-protein association, we focused the RIF generation process on non-polar sites in specific surface regions of interest: for example in the case of inhibitor design, interaction sites with biological partners. The RIF approach improves upon previous discrete interaction-sampling approaches^5^ by reducing algorithmic complexity from O(N) or O(N^2^) to O(1) with respect to the number of sidechain-target interactions considered, allowing for billions, rather than thousands, of potential interfaces to be considered.

For docking against the rotamer interaction field, it is desirable to have a very large set of protein scaffold options, as the chance that any one scaffold can house many interactions is small. The structure models of these scaffolds must be quite accurate so that the positioning is correct. Using fragment assembly^7^, piecewise fragment assembly^8^, and helical extension^9^, we designed a large set of miniproteins ranging in length from 50 to 65 amino acids containing larger hydrophobic cores than previous miniprotein scaffold libraries^1^, which makes the protein more stable and more tolerant to introduction of the designed binding surfaces. 84,690 scaffolds spanning 5 different topologies with structural metrics predictive of folding were encoded in large oligonucleotide arrays and 34,507 were found to be stable using a high-throughput proteolysis based protein stability assay^10^.

We experimented with several approaches for docking these stable scaffolds against the target structure rotamer interaction field, balancing overall shape complementarity with maximizing specific rotamer interactions. The most robust results were obtained using direct low resolution shape matching^11^ followed by grid based refinement of the rigid body orientation in the RIF (RIFDock). This resulted in better Rosetta binding energies (ddGs) and packing (contact molecular surface, see below) after sequence design than shape matching alone with PatchDock (**Fig. 1b red and green**), and more extensive non polar interaction with the target than hierarchical search without PatchDock shape matching ^6^ (**Extended Data Fig. 1**).

Because of the loss in resolution in the hashing used to build the RIF, and the necessarily approximate accounting for interactions between sidechains (see methods), we found that evaluation of the RIF solutions is considerably enhanced by full combinatorial optimization using the Rosetta forcefield, allowing the target sidechains to repack and the scaffold backbone to relax. Full combinatorial sequence optimization is quite CPU intensive, however, and to enable rapid screening through millions of alternative backbone placements, we developed a rapid pre-screening method using Rosetta to identify promising RIF docks. We found that including only hydrophobic amino acids, using a reduced set of rotamers than in standard Rosetta design calculations, and a more rapidly computable energy function sped design more than 10-fold while retaining a strong correlation with results after full sequence design (next paragraph); this pre-screen (referred to as the “Predictor” below) substantially improved the binding energies and shape complementarity of the final designs as far more RIF solutions could be processed (**Extended Data Fig. 2)**.

We observed that application of standard Rosetta design to the set of filtered docks in some cases resulted in models with buried unsatisfied polar groups and other suboptimal properties. To overcome these limitations, we developed a combinatorial sequence design protocol that maximizes shape and chemical complementarity with the target while avoiding buried polar atoms. Sequence compatibility with the scaffold monomer structure was increased using a structure based sequence profile^12^, the cross-interface interactions were upweighted during the Monte Carlo-based sequence design stage to maximize the contacts between the binder and the target (ProteinProteinInterfaceUpweighter; see Methods), and rotamers containing buried unsatisfiable polar atoms were eliminated prior to packing and buried unsatisfied polar atoms penalized by a pair-wise decomposable pseudo-energy term^13^. This protocol yielded amino acid sequences more strongly predicted to fold to the designed structure (**Extended Data Fig. 3a**) and to bind the target (**Extended Data Fig. 3b**) than standard Rosetta interface design.

In the course of developing the overall binder design pipeline, we noticed upon inspection that even designs with favorable Rosetta binding free energies, large changes in Solvent Accessible Surface Area (SASA) upon binding, and high shape complementarity (SC) often lacked dense packing and interactions involving several secondary structural elements. We developed a quantitative measure of packing quality in closer accord with visual assessment -- the contact molecular surface (see methods) -- which balances interface complementarity and size in a manner that explicitly penalizes poor packing. We used this metric to help select designs at both the rapid Predictor stage and after full sequence optimization (see Methods).

The space sampled by the search over structure and sequence space is enormous: tens of thousands of possible protein backbones × nearly one billion possible disembodied sidechain interactions per target × 10^16^ interface sequences per scaffold placement. Sampling of spaces of this size is necessarily incomplete, and many of the designs at this stage contained buried unsatisfied polar atoms (only rotamers that cannot make hydrogen bonds in any context are excluded at the packing stage) and cavities. To generate improved designs, we intensified the search around the best of the designed interfaces. We developed a resampling protocol which extracts all the secondary structural motifs making good contacts with the target protein from the first “broad search” designs, clusters these motifs based on their backbone coordinates and rigid body placements, and then selects the binding motif in each cluster with the best per-position weighted Rosetta binding energy; around 2,000 motifs were selected for each target. These motifs, which are privileged because they contain a much greater density of favorable side chain interactions with the target than the rest of the designs, were then used to guide another round of docking and design. Scaffolds from the library were superimposed on the privileged motifs, the favorable-interacting motif residues transferred to the scaffold, and the remainder of the scaffold sequence optimized to make further interactions with the target, allowing backbone flexibility to increase shape complementarity with the target (**Fig. 1a**). Interface metrics for the designs based on the resampling protocol were considerably improved relative to those of the designs from the broad searching stage (**Fig. 1b**). The designs with the most favorable protein folding and protein interface metrics from both the broad searching and resampling stages were selected for experimental validation.

### Experimental testing

Previous protein binder design approaches have been tested on only one or two targets, which limits assessment of their generality. To robustly test our new binder design pipeline, we selected thirteen native proteins of considerable current interest spanning a wide range of shapes and biological functions. These proteins fall into two classes: first, human cell surface or extracellular proteins involved in signaling, for which binders could have utility as probes of biological mechanism and potentially as therapeutics (Tropomyosin receptor kinase A (TrkA)^14^, Fibroblast growth factor receptor 2 (FGFR2)^15^, Epidermal growth factor receptor (EGFR)^16^, Platelet-derived growth factor receptor (PDGFR)^17^, Insulin receptor (InsulinR)^18^, Insulin-like growth factor 1 receptor (IGF1R)^19^, Angiopoietin-1 receptor (Tie2)^20^, Interleukin-7 receptor alpha (IL-7Rα)^21^, CD3 delta chain (CD3δ)^22^, Transforming growth factor beta (TGF-β)^23^); and second, pathogen surface proteins for which binding proteins could have therapeutic utility (Influenza A H3 hemagglutinin (H3)^24^, VirB8-like protein from Rickettsia typhi (VirB8)^25^, and the SARS-CoV-2 coronavirus spike protein) (**Fig. 2a**). For each target, we selected one or two regions to direct binders against for maximal biological utility and for potential downstream therapeutic potential. These regions span a wide range of surface properties, with diverse shape and chemical characteristics (**Fig. 2a and Extended Data Fig. 4**).

**Figure 2:**
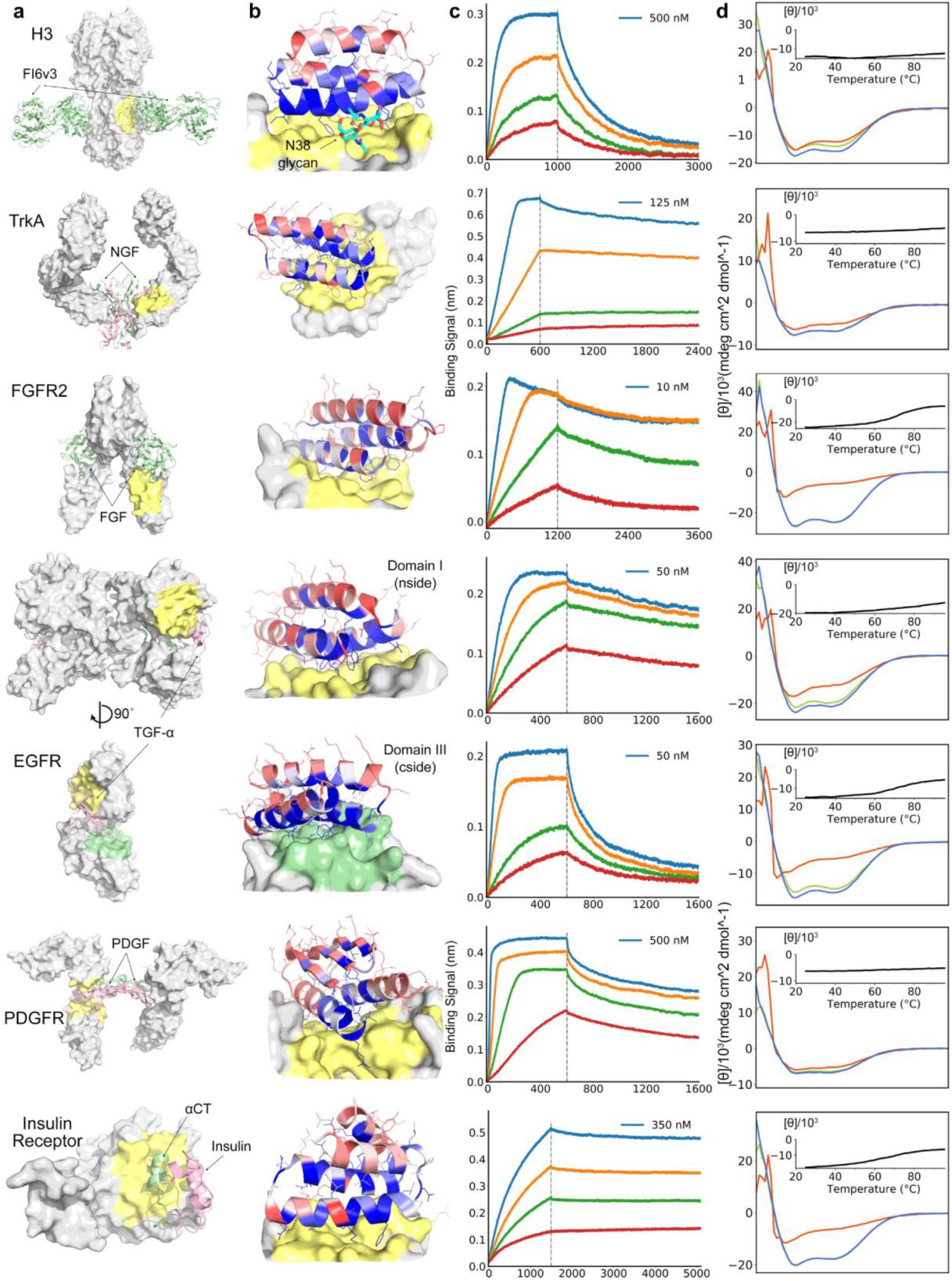
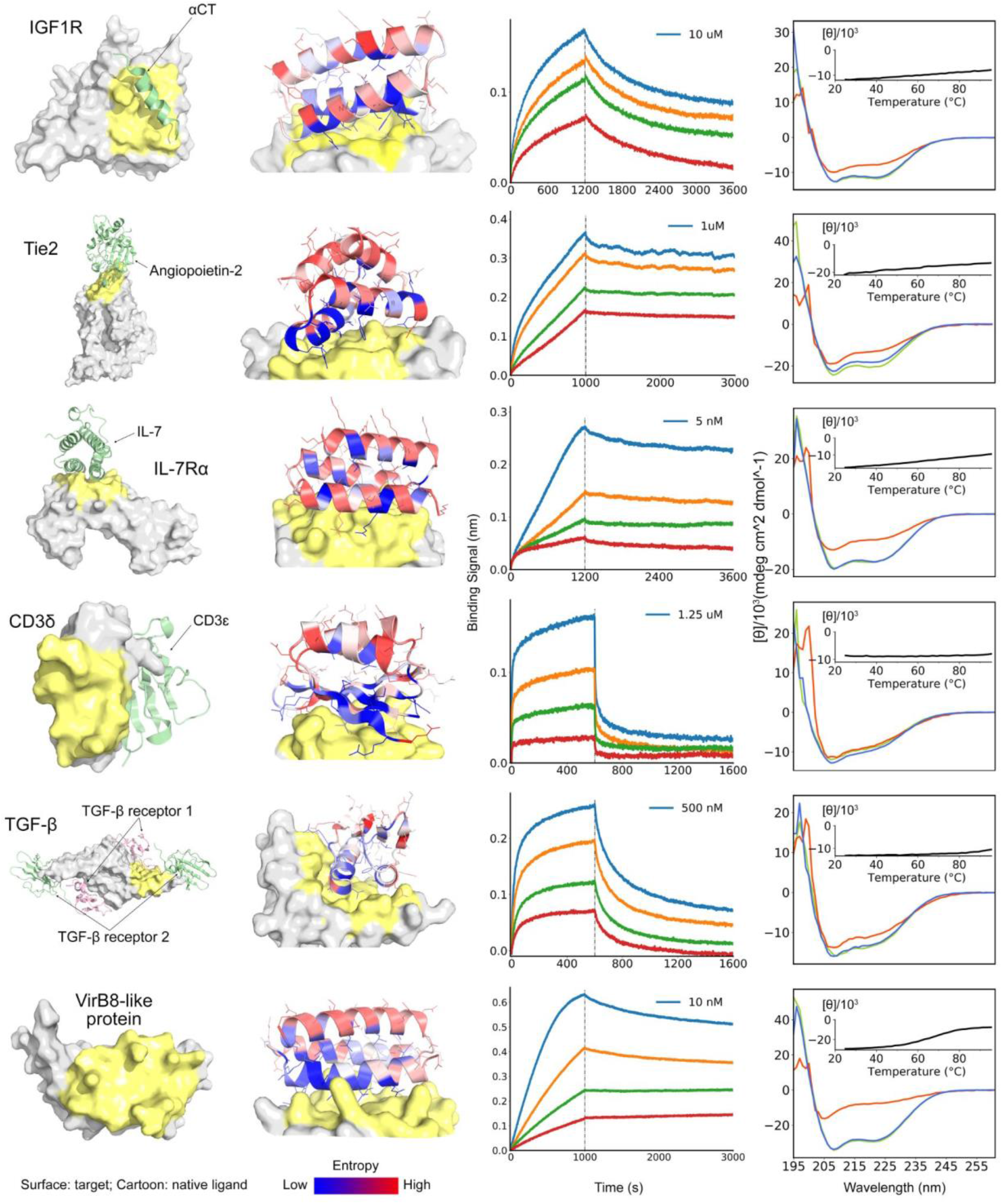
De novo design of miniprotein binders to 13 target sites. **a**, Naturally occurring target protein structures shown in surface representation, with known interacting partners f in cartoons where available. Regions targeted for binder design or in pale yellow or green; the remainder of the target surface is in grey. See (**Extended Data Figure 4**) for the zoomed in views of the selected targeting regions. The PDB ID codes are 3ZTJ (H3), 2IFG (TrkA), 1DJS (FGFR2), 1MOX (EGFR), 3MJG (PDGFR), 4OGA (InsulinR), 5U8R (IGF1R), 2GY7 (Tie2), 3DI3 (IL-7Rα), 1XIW (CD3δ), 3KFD (TGF-β) and 4O3V (VirB8). **b**, Computational models of designed complexes colored by site saturation mutagensis results. Designed binding proteins (cartoons) are colored by positional Shannon entropy, with blue indicating positions of low entropy (conserved) and red those of high entropy (not conserved); target surface is in grey and yellow. The core residues and binding interface residues are more conserved than the non-interface surface positions, consistent with the computational models. Full SSM maps over all positions of all the de novo designs are provided in (**Supplementary file/Extended Data Fig. 18)**. **c**, Biolayer interferometry characterization of binding of optimized designs to the corresponding targets. Two-fold serial dilutions were tested for each binder and the highest concentration is labeled. For H3, TrkA, FGFR2, EGFR, PDGFR, IL-7Rα, CD3δ, TGF-β and VirB8, the biotinylated target proteins were loaded onto the Streptavidin (SA) biosensors, and incubated with miniprotein binders in solution to measure association and dissociation. For IGF1R and Tie2, MBP- (maltose binding protein) tagged miniprotein binders were used as the analytes. For InsulinR, the miniprotein binder was immobilized onto the Amine Reactive Second-Generation (AR2G) Biosensors and the insulin receptor was used as the analyte. **d**, Circular dichroism spectra at different temperatures (green: 25 °C, red: 95 °C, blue: 95 °C followed by 25 °C) and (insert) CD signal at 222-nm wavelength as a function of temperature for the optimized designs.

Using the above protocol, we designed 15,000-100,000 binders for each of thirteen target sites on the twelve native proteins (see Methods; we chose two sites on the EGF receptor). Synthetic oligonucleotides (230bp) encoding the 50-65 residue designs were cloned into a yeast surface expression vector, the designs were displayed on the surface of yeast, and those which bind their target enriched by several rounds of fluorescence-activated cell sorting using fluorescently labelled target proteins. The starting and enriched populations were deep sequenced, and the fraction of each design after each sort was determined by comparing the frequency of the design in the parent and child pools. From multiple sorts at different target protein concentrations, we determined, as a proxy for binding Kd’s, the midpoint concentration (SC50) in the binding transitions for each design in the library (**Extended Data Table 1 and Supplementary Methods**).

To assess whether the top enriched designs for each target fold and bind as in the corresponding computational design models, and to investigate the sequence dependence of folding and binding, we generated high resolution footprints of the binding surface by sorting site saturation mutagenesis libraries (SSMs) in which every residue was substituted with each of the 20 amino acids one at a time. For the majority, but not all, enriched designs, substitutions at the binding interface and in the protein core were less tolerated than substitutions at non-interface surface positions (**Fig. 2b, Extended Data Fig. 20 & Extended Data Fig. 5**), and all the cysteines were highly conserved in designs containing disulfides. The effects of each mutation on both binding energy and monomer stability were estimated using Rosetta design calculations, and a reasonable correlation was found between the predicted and experimentally determined effect of mutations (**Extended Data Fig. 6**). In almost all cases, a small number of substitutions were found to increase apparent binding affinity, and we generated libraries combining 5-15 of these and sorted for binding under increasingly stringent (lower target concentrations) conditions. Many of these affinity-enhancing substitutions were mutations to tyrosine (**Extended Data Fig. 7**), consistent with the high relative frequency of tyrosine in natural protein interfaces^26^. The set of affinity increasing substitutions provide valuable information for improving the approach as these substitutions ideally would have been identified in the computational sequence design calculations (see discussion below).

We expressed the highest affinity combinatorially-optimized binders for each target in *E.coli* for more detailed structural and functional characterization. All of the designs were in the soluble fraction, and could be readily purified by nickel-NTA chromatography. All had circular dichroism spectra consistent with the design model, and most (9 out of 13) were stable at 95 °C (**Fig. 2d**). The binding affinities for the targets were assessed by biolayer interferometry, and found to range from 300 pM to 900 nM (**Fig. 2c and Extended Data Table 2**). The sequence mapping data report on the residues on the design critical for binding, but only weakly on the region of the target bound. We investigated this using a combination of binding competition experiments, biological assays, and structural characterization of the complexes. For the nine targets for which these were available, this characterization suggested binding modes consistent with the design models, as described in the following paragraphs.

### Host protein targets involved in signaling

The receptor tyrosine kinases TrkA, FGFR2, PDGFR, EGFR, InsulinR, IGF1R and Tie2 are key regulators of cellular processes and are involved in the development and progression of many types of cancer^27^. We designed binders targeting the native ligand binding sites for PDGFR, EGFR (on both domain I and domain III, the binders are referred to as EGFRn_mb and EGFRc_mb respectively), InsulinR, IGF1R and Tie2, and targeting surface regions proximal to the native ligand binding sites for TrkA and FGFR2 (**Fig. 2a** and see methods for criteria). We obtained binders to all eight target sites; the binding affinities of the optimized designs ranged from ~1nM or better for TrkA and FGFR2, to 860nM for IGF1R. Competition experiments with nerve growth factor (NGF), Platelet Derived Growth Factor-BB (PDGF-BB), insulin, insulin growth factor-1 (IGF-1) and Angiopoietin 1 (Ang1) on yeast suggest that the binders for TrkA, PDGFR, InsulinR, IGF1R and Tie2 bind to the targeted sites (**Extended Data Fig. 8**), consistent with the computational design models. The receptor tyrosine kinase binders as monomers are all expected to be antagonists, and we tested the effect on signaling through TrkA, FGFR2 and EGFR of the cognate binders on cells in culture. Strong inhibition of signaling by the native agonists was observed in all three cases (**Fig. 3a-c, Extended Data Fig. 9 and Extended Data Fig. 10**).

**Figure 3:**
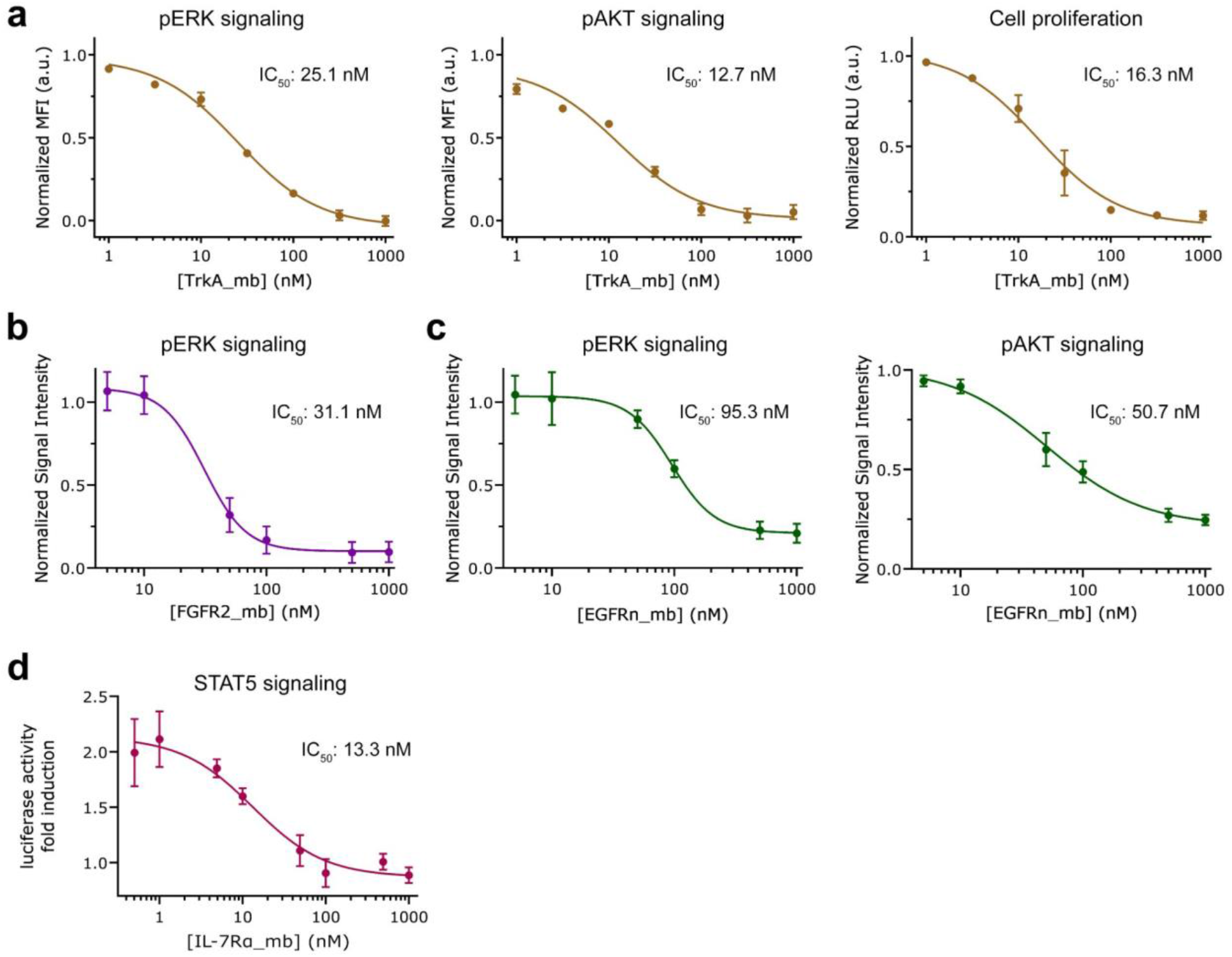
Inhibition of native signaling pathways by designed miniprotiens. **a**, Dose-dependent reduction in (left) pERK signaling, (middle) pAKT signaling and cell proliferation after 48 hrs (right) of TF-1 cells with increase in TrkA minibinder concentration. 8.0 ng/ml human beta-NGF was used for competition. Titration curves at different concentrations of NGF and the effects of the miniprotein binders on cell viability are in **Extended Data Fig. 9. b**, Dose-dependent reduction pERK signaling elicited by 0.75 nM bFGF in HUVECs with increasing FGFR2 minibinder concentration. **c**, Dose-dependent reduction in (left) pERK signaling, and (right) pAKT signaling elicited by 1nM EGF in HUVECs with increase in EGFR n-side minibinder concentration. See **Extended Data Fig. 10** and methods for the experimental details. **d**, Reduction in STAT5 activity induced by 50 pM of hIL-7 in HEK293T cells in the presence of increased hIL-7Rα minibinder concentrations. IC50 was calculated using a four-parameter-logistic

Binding of IL-7 to the IL-7α receptor subunit leads to recruitment of the γc receptor, forming a tripartite cytokine-receptor complex crucial to several signaling cascades leading to the development and homeostasis of T and B cells^28^. We obtained a picomolar affinity binder for IL-7Rα targeting the IL-7 binding site, and found that it blocks STAT5 signaling induced by IL-7 (**Fig. 3d**). We also obtained binders to CD3δ, one of the subunits of the T-cell receptor, and the signaling molecule TGF-β, which play critical roles in immune cell development and activation (**Fig. 2**).

### Pathogen target proteins

Hemagglutinin (HA) is the main target for influenza A virus vaccine and drug development, and it can be genetically classified into two main subgroups, group 1 and group 2^29,30^. The HA stem region is an attractive therapeutic epitope, as it is highly conserved across all the influenza A subtypes and targeting this region can block the low pH-induced conformational rearrangements associated with membrane fusion, which is essential for virus infection^31,32^. Neutralizing antibodies targeting the stem region of group 2 HA have been identified through screening of large B-cell libraries after vaccination or infection that neutralize both group 1 and group 2 influenza A viruses^33,34^. Protein ^1,5^, peptide^35^ and small molecule inhibitors^36^ have been designed to bind to the stem region of group 1 HA and neutralize the influenza A viruses, but none recognize the group 2 HA. However, the design of small proteins or peptides that can bind and neutralize both group 1 HA and group 2 HA has been challenging due to three differences between the group 1 HA and the group 2 HA: first, the group 2 HA stem region is more hydrophilic, containing more polar residues, second, in group 2 HA, Trp21 adopts a configuration roughly perpendicular to the surface of the targeting groove, which makes the targeted groove much shallower and less hydrophobic, and third, the group 2 HA is glycosylated at Asn38 with the carbohydrate side chains covering the hydrophobic groove (**Extended Data Fig. 11**). We used our new method to design binders to H3 HA (A/Hong Kong/1/1968), the main pandemic subtype of group 2 influenza virus, and obtained a binder with an affinity of 320 nM to the wild type H3 (**Fig 2**) and 28nM to the deglycosylated H3 variant (N38D) (**Extended Data Fig. 12a**); the reduction in affinity is likely due to the entropy loss of the glycan upon binding and/or the steric clash with the glycan. The binder also binds to H1 HA (A/Puerto Rico/8/1934) which belongs to the main pandemic subtype of group 1 influenza virus (**Extended Data Fig. 12b**); the binding with both H1 and H3 is competed by the stem region binding neutralizing antibody FI6v3^33^ on the yeast surface (**Extended Data Fig. 12c,d**), suggesting that the binder binds the hemagglutinin at the targeted site. We also designed binders to the prokaryotic pathogen protein VirB8 which belongs to the type IV secretion system of *Rickettsia typhi*, which is the causative agent of murine typhus^25^. We selected the surface region composed of the second and the third helices of VirB8, and obtained binders with 500 pM affinity (**Fig. 2**).

With the outbreak of the SARS-CoV-2 coronavirus pandemic we applied our method to design miniproteins targeting the receptor binding domain of the SARS-CoV-2 Spike protein near the ACE2 binding site to block receptor engagement. Due to the pressing need for coronavirus therapeutics, we recently described the results of these efforts^37^ ahead of those described in this manuscript; As in the case of FGFR2, IL-7Rα and VirB8, the method yielded picomolar binders, which are among the most potent compounds known to inhibit the virus in cell culture (IC50 0.15ng/ml) and subsequent animal experiments have shown that they provide potent protection against the virus in vivo^38^. The modular nature of the miniprotein binders enables their rapid integration into designed diagnostic biosensors for both influenza and SARS-CoV-2 binders^39^.

The designed binding proteins are all very small proteins (<65 amino acids), and many are 3-helix bundles. To evaluate their target specificity, we tested the highest affinity binder to each target for binding to all other targets. There was very little cross reactivity (**Fig. 4a)**, likely due to their quite diverse surface shapes and electrostatic properties (**Fig. 4b**). Consistent with previous observations with affibodies^40^, this suggests that a wide variety of binding specificities can be encoded in simple helical bundles; in our approach, scaffolds are customized for each target, so the specificity arises both from the set of sidechains at the binding interface, and the overall shape of the interface itself.

**Figure 4:**
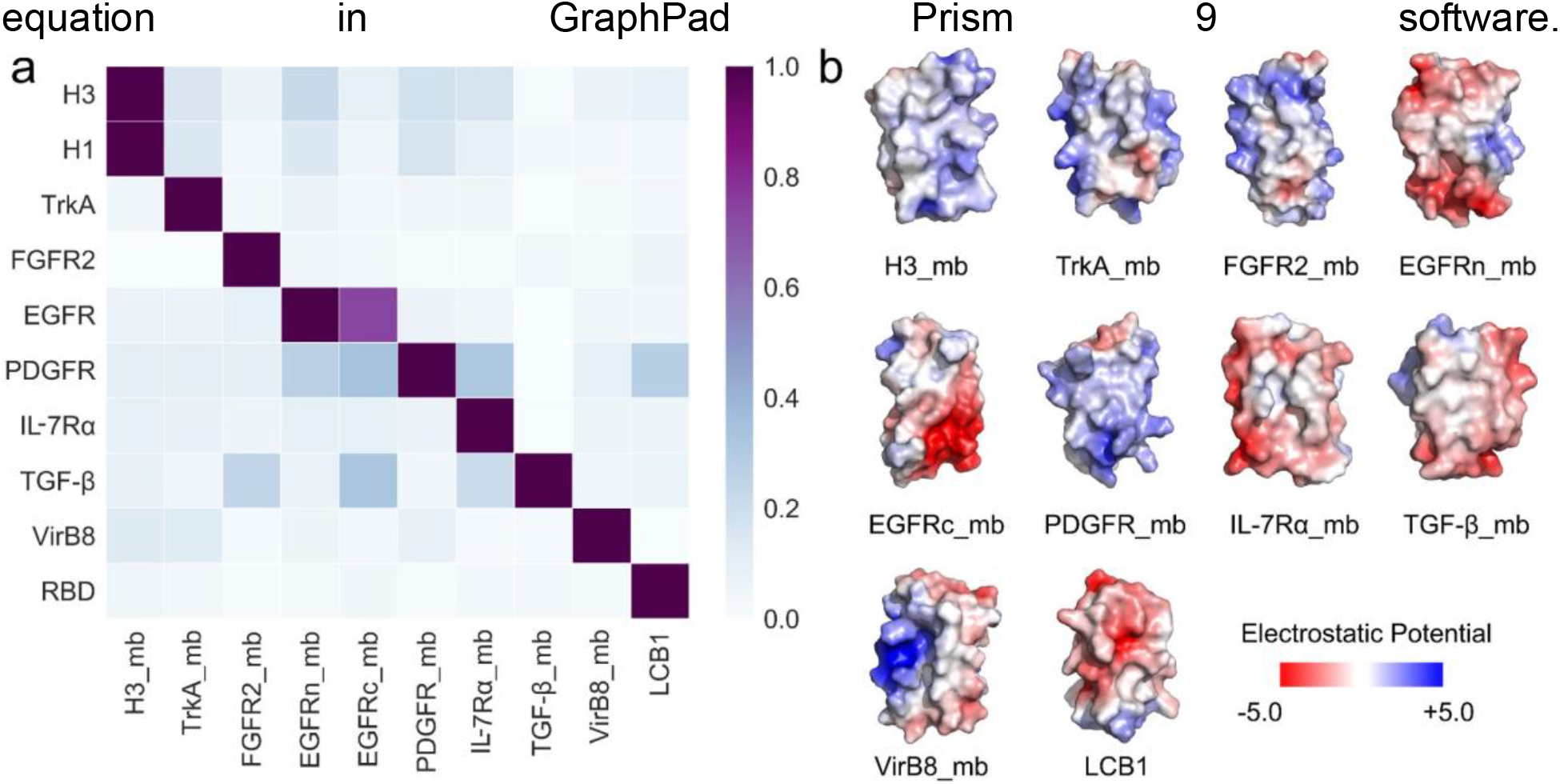
Designed binders have high target specificity. To assess the cross reactivity of each miniprotein binder with each target protein, The biotinylated target proteins were loaded onto biolayer interferometry SA sensors, allowed to equilibrate, and baseline signal set to zero. The BLI tips were then placed into 100 nM binder solution for 300 seconds, washed with buffer, and dissociation was monitored for an additional 600 seconds. Heatmap shows the maximum response signal for each binder-target pair normalized by the maximum response signal of the cognate designed binder-target pair. **b**, Surface shape and electrostatic potential (generated with the APBS Electrostatics plugin in Pymol; red positive, blue, negative) of the designed binding interfaces.

### High-resolution structural validation

High resolution structures are critical for evaluating the accuracy of computational protein designs. We succeeded in obtaining crystal structures of the unbound miniprotein binders for FGFR2 and IL-7Rα, as well as the co-crystal structures of the miniprotein binders of H3, TrkA, FGFR2 and IL-7Rα in complex with their targets (**Extended Data Table 3**). The H3 binder binds to the shallow groove of the stem region of HK68/H3 HA in the crystal structure as designed; the Cα root-mean-square deviation (rmsd) over the entire miniprotein binder is 1.42 Å using the HA as the alignment reference (**Fig. 5a and Extended Data Fig. 13**). The binder makes extensive hydrophobic interactions with HA and almost all of the designed interface side chain configurations are recapticulated in the crystal structure (**Fig. 5a**). There is a clear reorientation of the oligosaccharide at Asn38 compared with the unbound HK68/H3 structure (**Fig. 5a and Extended Data Fig. 11**; this has been observed in HK68/H3 structures bound with stem region neutralizing antibodies ^33,34^), consistent with the higher binding affinity for a deglycosylated variant (N38D) than for wild type H3 HA (A/Hong Kong/1/1968) in BLI assays (**Fig. 2 and Extended Data Fig. 12**). The crystal structure of the TrkA binder in complex with TrkA was very close to the design model (**Fig. 5b**). After aligning the crystal structure and design model on TrkA, the Cα rmsd over the entire miniprotein binder is 2.41 Å, and over the two interfacial binding helices 1.20 Å. The crystal structures of the FGFR2 binder by itself (**Extended Data Fig. 14a**) and in complex with the third Ig-like domain of FGFR4 (**Fig. 5c**) match the design models with near atomic accuracy, with Cα rmds of 0.58 Å for the binder alone and 1.87 Å over the entire complex. The TrkA binder and the FGFR2 binder bind to the curved sheet side of the ligand binding domain of TrkA and FGFR4 with extensive hydrophobic and polar interactions, and most of the key hydrophobic interactions as well as the primarily polar interactions in the computational design models are largely recapitulated in the crystal structures (**Fig 5b,c**). The binding interface partially overlaps with the native ligand binding sites of nerve growth factor (NGF) and fibroblast growth factor (FGF), however, the detailed sidechain interactions are entirely different in the designed and native complexes (**Extended Data Fig. 15a,b**). For IL-7Rα, the crystal structure of the monomer is close to that of the design, with a Cα rmsd of 0.63 Å (**Extended Data Fig. 14b**) and the co-crystal structure with IL-7Rα also matches with the design model closely, with a Cα rmsd of 2.2 Å using IL-7Rα as the reference (**Fig 5d**). Both the de novo IL-7Rα binder and the native IL-7 use two helices to bind with IL-7Rα, but the binding orientations are totally different (**Extended Data Fig. 15c**). Further highlighting the accuracy of the protein interface design method, the cryoEM structures of the SARS-CoV-2 binders LCB1 and LCB3 in complex with the virus are also nearly identical to the design models, with Cα rmsd of 1.27 Å and 1.9 Å respectively^37^ (**Fig. 5e**). While we were not able yet to solve structures for the remainder of the designs, the high resolution sequence footprinting (**Fig. 2b, Extended Data Fig. 20 & Extended Data Table 4**) and competition results suggest that the interfaces involve both the designed residues and the intended regions on the target. The very close agreement between the experimentally determined structures and the original design models suggests that the substitutions required to achieve high affinity play relatively subtle roles in tuning interface energetics; the overall structure of the complex, including the structure of the monomer binders and the detailed target binding mode, are determined by the computational design procedure.

**Figure 5:**
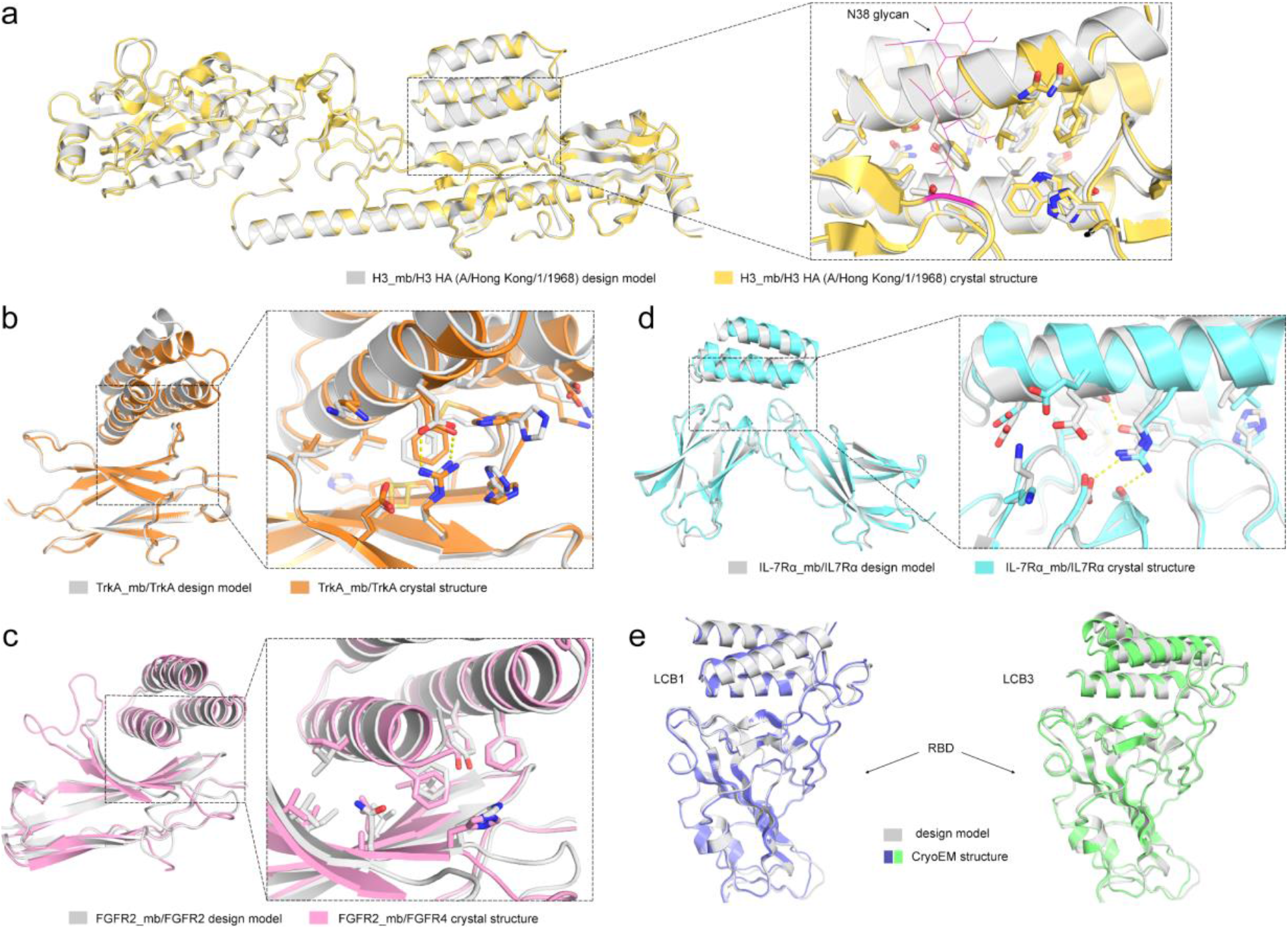
High-resolution structures of miniprotein binders in complex with target proteins are very close to computational design models. **(a-d).** (left) Superimposition of computational design model (silver) on experimentally determined crystal structure. (right) Zoom-in view of designed interface, with interacting side chains as sticks. **a.** H3, **b.** TrkA, c. FGFR2, **d.** IL-7Rα. **e**, Superimposition of the computational design model and refined cryo-EM structures of LCB1 (left) and LCB3 (right) bound to receptor binding domain of SARS-CoV-2 spike protein (design models are in gray and cryoEM structures are in pale blue and green).

### Determinants of design success

For our de novo design strategy to be successful, we must encode in the ~60 residue designed sequences both information on the folded monomer structures, and on the target binding interfaces: designs which do not fold to the correct structure, or which fold to the intended structures but do not bind to the target will fail. To assess the accuracy with which the monomer structure must be designed, we carried out an additional calculation and experiment for the IL-7Rα target. Large numbers of scaffolds were superimposed onto 11 interface helical binding motifs identified in the first broad design search, and sequence design was carried out as described above. There was a strong correlation between the extent of binding and the RMSD to the binding motif (**Extended Data Fig. 16**), suggesting that designed backbones must be quite accurate to achieve binding. To assess the determinants of binding of the designed interfaces, assuming that the designs fold to the intended monomer structures, we took advantage of the large data set (810,000 binder designs and 240,000 single mutants) generated in this study. Across all targets, there was a strong correlation between success rate and the hydrophobicity of the targeted region (**Extended Data Fig. 17**), and designs observed experimentally to bind their targets tended to have stronger predicted binding energy, and larger contact molecular surfaces (**Extended Data Fig. 18**). As found previously for design of protein stability^10^, iterative design-build-test cycles in which the design method is updated at each iteration to incorporate feedback from the previous design round should lead to systematic improvement in the design methodology and success rate.

## Conclusions

Our success in designing nM affinity binders for 14 target sites demonstrates that binding proteins can be designed de novo using only information on the structure of the target protein, without need for prior information on binding hotspots or fragments from structures of complexes with binding partners. The success also suggests that our design pipeline provides a quite general solution to the de novo protein interface design problem that goes far beyond previously described methods. However, there is still considerable room for improvement. Only a small fraction of designs bind, and in almost all cases, the best of these require a few additional substitutions to achieve high affinity binding (**Extended Data Table 2**). Furthermore, the design of binders to highly polar target sites remains a considerable challenge-the sites targeted here all contain at least four hydrophobic residues. The datasets generated in this work -- both the information on binders versus non binders, and the feedback on the effects of individual point mutants on binding -- should help guide the development of methods for designing high affinity binders directly from the computer with no need for iterative experimental optimization. More generally, the de novo binder design method and the large data set generated here provide a starting point for investigating the fundamental physical chemistry of protein-protein interactions, and for developing and assessing computational models of protein-protein interactions.

This work is a major step forward towards the longer range goal of direct computational design of high affinity binders starting from structural information alone. We expect the binders created here, and new ones created with the method moving forward, will find wide utility as signaling pathway antagonists as monomeric proteins and as tunable agonists when rigidly scaffolded in multimeric formats, and in diagnostics and therapeutics for pathogenic disease. Unlike antibodies, the designed proteins can be expressed solubly in *E. coli* at high levels and are thermostable, and hence could form the basis for a next generation of lower cost protein therapeutics. More generally, the ability to rapidly and robustly design high affinity binders to arbitrary protein targets could transform the many areas of biotechnology and medicine that rely on affinity reagents.

## Acknowledgements

This work was supported by DARPA Synergistic Discovery and Design (SD2) HR0011835403 contract FA8750-17-C-0219 (L.C., B.C., S.H., D.B.), The Audacious Project at the Institute for Protein Design (L.K), the Open Philanthropy Project Improving Protein Design Fund (B.C., D.B.), funding from Eric and Wendy Schmidt by recommendation of the Schmidt Futures program (I.G., L.M.), an Azure computing resource gift for COVID-19 research provided by Microsoft (L.C., B.C.), the National Institute of Allergy and Infectious Diseases (HHSN272201700059C, D.B., B.H., L.S.; NIH R01 AI140245 to E.M.S.; NIH R01 AI150855 to I.A.W.), the National Institute on Aging (R01AG063845, B.H., D.B.), the Defense Threat Reduction Agency (HDTRA1-16-C-0029, D.B. E-M.S.), The Donald and Jo Anne Petersen Endowment for Accelerating Advancements in Alzheimer’s Disease Research (N.B.), a gift from Gates Ventures (M.D.), The Human Frontier Science Program (A.Y.) and The Howard Hughes Medical Research Institute (K.M.J, K.C.G., D.B.). Use of SSRL at Stanford Linear Accelerator Center (SLAC) National Accelerator Laboratory is supported by the US Department of Energy Office of Science, Office of Basic Energy Sciences under contract DE-AC02-76SF00515. The SSRL Structural Molecular Biology Program is supported by the Department of Energy, Office of Biological and Environmental Research and the National Institutes of Health, National Institute of General Medical Sciences (including P41GM103393). A part of this work is based upon research conducted at the Northeastern Collaborative Access Team beamlines, which are funded by the National Institute of General Medical Sciences from the National Institutes of Health (P30 GM124165). The Eiger 16M detector on the 24-ID-E beam line is funded by a NIH-ORIP HEI grant (S10OD021527). STRW was supported by the CCR intramural research program of NCI-NIH. GM/CA at the Advanced Photon Source at Argonne National Laboratory has been funded by the National Cancer Institute (ACB-12002) and the National Institute of General Medical Sciences (AGM-12006, P30GM138396). This research used resources of the Advanced Photon Source, a U.S. Department of Energy (DOE) Office of Science User Facility operated for the DOE Office of Science by Argonne National Laboratory under Contract No. DE-AC02-06CH11357. The Eiger 16M detector at GM/CA-XSD was funded by NIH grant S10 OD012289. We thank the staff of beamline ID23-2 (ESRF) for technical support and beamtime allocation. S.N.S. acknowledges research support from Research Foundation Flanders (grants G0C2214N and G0E1516N), and the Hercules Foundation (no. AUGE-11-029). S.N.S. is a principal investigator of the VIB (Belgium).

We thank George Ueda for kindly providing the Ang1 protein for the TrkA competition assay and Deborah H. Fuller for kindly providing the FI6v3 antibody for the HA competition assay. We thank Yong-Jun Park, Alexandra Walls, and David Veesler for their collaborative research and cryoEM structure determination for minibinders targeting SARS-CoV-2 Spike. We would also like to thank Kandise Van Wormer and Austin Curtis Smith for their tremendous laboratory support during COVID-19.

